# A simulation-based evaluation of STRUCTURE software for exploring the introduction routes of invasive species

**DOI:** 10.1101/094029

**Authors:** Eric Lombaert, Thomas Guillemaud, Emeline Deleury

**Author notes:** Corresponding author: Eric Lombaert, INRA, UMR 1301 IBSV (INRA / Université de Nice Sophia Antipolis / CNRS). 400 Route des, Chappes. BP 167 - 06903 Sophia Antipolis cedex. FRANCE.

## Abstract

Population genetic methods are widely used to retrace the introduction routes of invasive species. The unsupervised Bayesian clustering algorithm implemented in STRUCTURE is amongst the most frequently use of these methods, but its ability to provide reliable information about introduction routes has never been assessed. We used computer simulations of microsatellite datasets to evaluate the extent to which the clustering results provided by STRUCTURE were misleading for the inference of introduction routes. We focused on the simple case of an invasion scenario involving one native population and two independently introduced populations, because it is the sole scenario with two introduced populations that can be rejected when obtaining a particular clustering with a STRUCTURE analysis at *K* = 2 (two clusters). Results were classified as “misleading” or “non-misleading”. We then investigated the influence of two demographic parameters (effective size and bottleneck severity) and different numbers of loci on the type and frequency of misleading results. We showed that misleading STRUCTURE results were obtained for 10% of our simulated datasets and at a frequency of up to 37% for some combinations of parameters. Our results highlighted two different categories of misleading output. The first occurs in situations in which the native population has a low level of diversity. In this case, the two introduced populations may be very similar, despite their independent introduction histories. The second category results from convergence issues in STRUCTURE for *K* = 2, with strong bottleneck severity and/or large numbers of loci resulting in high levels of differentiation between the three populations.

## Introduction

Retracing the introduction routes of invasive alien species is a prerequisite to accurately compare ancestral and derived populations to infer ecological and evolutionary processes which determine the invasion success. However, identification of the source of an introduced population is a complex task, because of the highly stochastic nature of the introduction process (Estoup and Guillemaud, 2010). Many population genetics methods and tools are now widely used to retrace the introduction routes of invasive species. This approach is somewhat risky, because the methods involved are often dependent on demographic and genetic equilibria, but invasions often involve demographic disequilibrium, through strong bottlenecks followed by rapid population growth, for example. Despite this limitation and the risks of using population genetics methods inappropriately in the specific context of biological invasions, only a few of these methods have been formally evaluated (e.g. Estoup and Guillemaud, 2010; Guillemaud *et al.*, 2010).

Among population genetics methods, unsupervised individual Bayesian clustering methods are widely used. The popularity of these methods is due to their ability to infer genetic structure correctly in many situations and their apparent simplicity (several “click-and-play” software suites are available). STRUCTURE (Pritchard *et al.*, 2000; Falush *et al.*, 2003; Hubisz *et al.*, 2009) is the most frequently used software for clustering, with more than 27,000 citations for the three references indicated above in Google Scholar in May 2017. STRUCTURE aims to sort individuals in an unsupervised way into *K* clusters (*K* being defined by the user), assuming Hardy-Weinberg/linkage equilibrium within clusters (Porras-Hurtado *et al.*, 2013). In theory, if *K* is set to the true number of population, samples belonging to the same population will be classified into the same cluster. More broadly, because knowing or inferring the true number of population is not always possible, samples belonging to the same cluster are at least considered as sharing a close evolutionary history. STRUCTURE is known to perform well in most cases, but it can be misleading in some situations, particularly in the presence of isolation by distance (Frantz *et al.*, 2009; Schwartz and McKelvey, 2009), clusters of very different sizes (Kalinowski, 2011; Puechmaille, 2016), family groups (Anderson and Dunham, 2008), or high proportions of missing data (Smith and Wang, 2014).

STRUCTURE and other software suites based on similar methods are frequently used in the context of introduction routes inferences (Estoup and Guillemaud, 2010; Lawson Handley *et al.*, 2011; Cristescu, 2015). In some cases, STRUCTURE is used directly to contrast models of invasion history, mainly for comparisons of scenarios involving either multiple independent introductions from a native population, or a single introduction from the native area followed by subsequent introduction(s) from this primary introduced area. In this context, exploring clustering patterns with only two genetic clusters (*K* = 2) is considered as informative. Indeed, one of the clustering patterns that can be obtained makes it possible to reject the hypothesis of independent introductions: if all samples from the invaded areas group together in one cluster, and all samples from the native area group in the other cluster, this allows rejecting the hypothesis of independent introductions and is considered to provide fairly conclusive evidence about a single introduction from the native area (Fig. 1). For example, Ascunce *et al.* (2011) explored the worldwide invasion history of the fire ant *Solenopsis invicta* with a total of 2,144 colonies sampled from 75 geographic locations, including 39 native (South America) and 36 invaded (USA, China, Australia) areas. They found that all samples from invasive populations clustered together when analyzing the data with STRUCTURE at *K* = 2 and concluded that only one introduction from the native area occurred. They then used approximate Bayesian computation to test whether the oldest invasive population in the USA was the source of all other invasive populations in distant areas. Similarly, Cordero *et al.* (2017) analyzed 378 individuals of the Manila clam *Ruditapes philippinarum* from 9 geographic locations, including 3 native (Asia) and 6 invaded (North America and Europe) areas. They found that STRUCTURE analyses at *K* = 2 grouped all samples of invasive populations into the same cluster. They concluded that a single native Asian introduction of the species into North America was very likely, and that North America then became the source of the European outbreak. Such use of STRUCTURE in the context of invasion biology is very common (e.g. Lachmuth *et al.*, 2010; Papura *et al.*, 2012; Robert *et al.*, 2012; Bolte *et al.*, 2013; Fontaine *et al.*, 2013; Sanz *et al.*, 2013; Zhang *et al.*, 2014; Yu *et al.*, 2014; Zhou *et al.*, 2015; Guillemaud *et al.*, 2015; Rewicz *et al.*, 2015; Dieni *et al.*, 2016; Zhu *et al.*, 2017). However, invasions frequently involve major demographic events, such as strong bottlenecks followed by genetic drift, which may significantly impair our ability to determine introduction routes correctly from a given STRUCTURE result. This may account for the contradictory outcomes sometimes obtained with different population genetics methods. For example, Mallez *et al.* (2015) found conflicting results when trying to infer the origin of the invasive Portuguese outbreak of the pinewood nematode *Bursaphelenchus xylophilus*: while *F*_ST_ values suggested a native North American origin, STRUCTURE suggested an origin from an oldest invasive population in Japan for these samples, because all invasive samples from Portugal and Japan belonged to one cluster and all native samples belonged to another cluster while analyzing *K* = 2 patterns.

**Figure 1:**
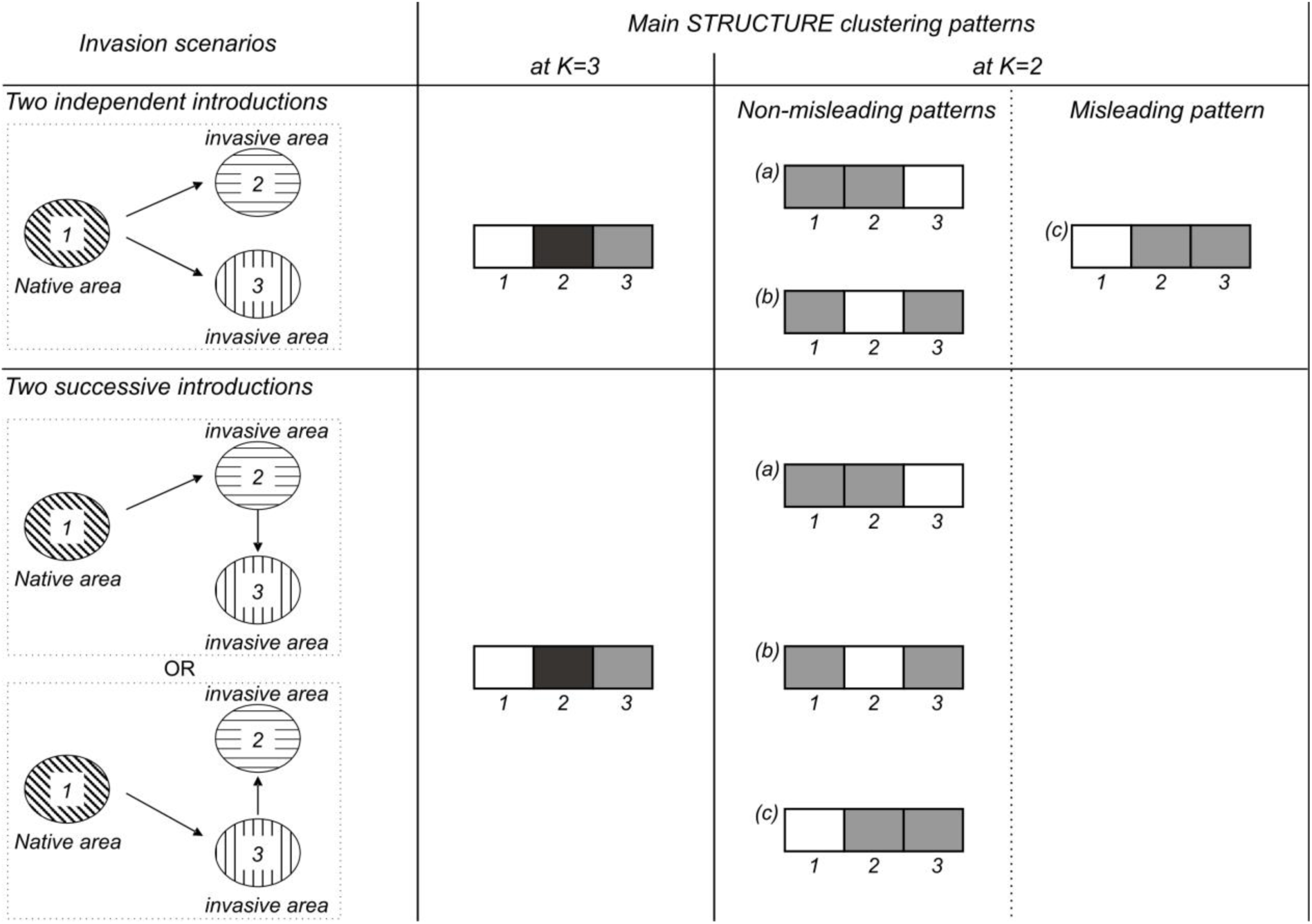
Schematic representations of the main STRUCTURE clustering patterns that can be obtained at *K* = 3 and *K* = 2 according to different invasion scenarios (either independent or successive) involving one native and two invasive populations. Other patterns with admixture are also possible but are not shown here because they are less informative in the context of invasion routes. At *K* = 3, with 3 samples, the same pattern (i.e. each sample constitutes a cluster) is likely to be found whatever the scenario, and thus no valuable information about the origin of introduced populations can be deduced. On the contrary, patterns obtained at *K* = 2 can be informative: whereas clustering patterns *a* and *b* can be obtained in both independent and successive scenario, pattern *c* should only be found if introductions are successive. If obtained, this pattern *c* would lead a STRUCTURE user to eliminate the independent scenario as a likely one.

In this study, we evaluated the risk of incorrect introduction route inferences based on STRUCTURE analyses, for the simple case of an invasion scenario involving one native population and two independently introduced populations. We chose to simulate this scenario because it is the sole one that can be rejected when obtaining a particular clustering with a STRUCTURE analysis at *K* = 2 (Fig. 1). We simulated a large number of microsatellite datasets drawn from populations of various effective sizes and bottleneck severities. STRUCTURE analyses were performed on these simulated datasets and the resulting clustering patterns at *K* = 2 were classified as “misleading” or “non-misleading”. We then explored the effect of demographic parameters on the likelihood of misleading patterns being obtained, to identify and predict the situations in which the use of STRUCTURE in a context of introduction routes inference may be risky.

## Methods

### Scenario description and data simulation

We chose to simulate a scenario with two independent introductions because it is the only one that can be rejected from a STRUCTURE analysis when considering two introduced populations and a native one (Fig. 1). We thus defined a simple historical scenario in which two invasive populations (populations 2 and 3) were independently founded 50 generations ago from the same native population (population 1). Both invasive populations were subject to a demographic bottleneck lasting 20 generations (Fig. 2a). The effective sizes of all three populations at equilibrium (*N*) and the effective number of founders of the two invasive populations during the bottlenecks (*NF*) could take different values: 10000, 1000, 100, 10 and 2 individuals, with *N* ≥ *NF.* Log_10_(*N*/*NF*) was considered to quantify bottleneck severity.

**Figure 2:**
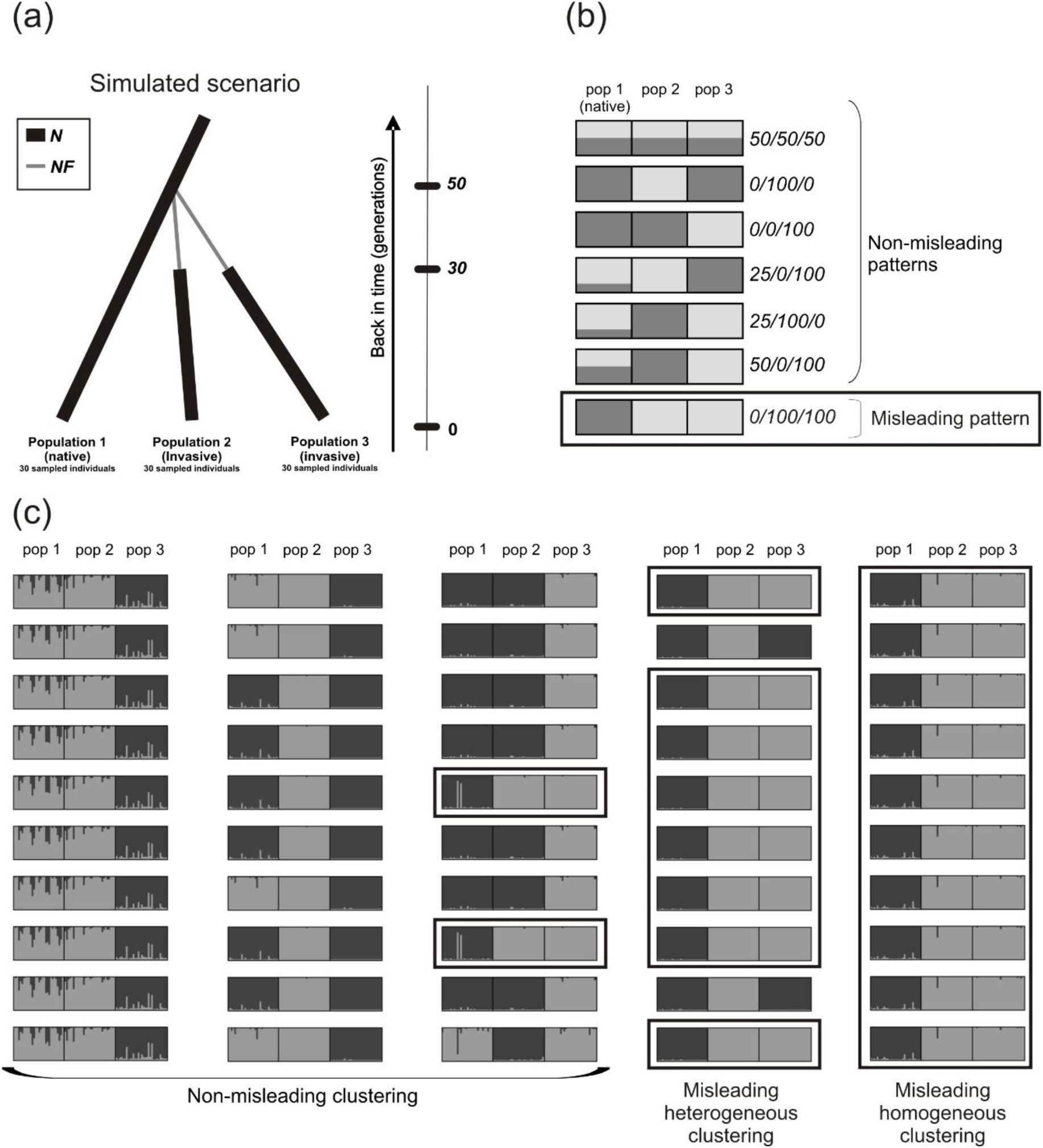
Simulated scenario and main observed STRUCTURE patterns at *K* = 2. (**a**) Graphical representation of the simulated scenario in which two invasive populations (populations 2 and 3) are independently derived from the native population (population 1). *N* is the effective size at equilibrium and *NF* is the effective number of founders during the bottlenecks. (**b**) Schematic representations of the main patterns obtained in the STRUCTURE runs for *K* = 2 and their associated summarized codes. The misleading pattern, inconsistent with the simulated scenario, is boxed. (**c**) Five examples of clusterings obtained over ten STRUCTURE runs for *K* = 2, and their associated classification. In this study, we focused on “misleading homogeneous clusterings” and “misleading heterogeneous clusterings”, in which “misleading patterns” were found in all ten runs or predominated, respectively, for a given dataset. Runs displaying the misleading pattern are boxed.

We used DIYABC version 2.0.4 software (Cornuet *et al.*, 2014) to generate 500 microsatellite multilocus genotype datasets for each of the 15 different combinations of *N* and *NF* values, through a coalescent process. For all datasets, a sample of 30 diploid individuals per population was simulated. We evaluated the effect of the number of loci on the analyses, by performing simulations with 10, 20 and 100 unlinked microsatellite markers. We used a generalized stepwise mutation model, with realistic values for all three parameters (Jarne and Lagoda, 1996; Estoup *et al.*, 2002): the mean mutation rate (set to 5×10^−4^), the mean parameter of the geometric distribution defining the number of microsatellite repeats gained or lost during mutation events (set to 0.22) and the mean mutation rate for single-nucleotide insertion/deletion (set to 10^−8^). In total, we simulated 22,500 datasets (15 sets of parameters × 500 datasets per set x 3 numbers of loci). We developed a pipeline with PERL scripts, available on request, to automate the processing of the datasets (simulation and subsequent STRUCTURE and post-STRUCTURE analyses).

### STRUCTURE analyses and misleading clustering

For each of the 22,500 simulated datasets, a Bayesian clustering analysis was performed in parallel, on a 120-nodes computer cluster, with STRUCTURE software version 2.3.4 (Pritchard *et al.*, 2000). We chose the admixture model with correlated allele frequencies. We used default values for all the other parameters. Each run consisted of a burn-in period of 10^5^ Markov chain Monte Carlo (MCMC) iterations, followed by 5×10^5^ MCMC iterations. This run length is considered to be long enough to obtain precise estimates of parameters (Pritchard *et al.*, 2010), but we also tried runs of double this length for some combinations of parameters with 100 loci. The results obtained were the same (data not shown). We carried out ten replicate runs for each dataset and each value of *K*, the number of genetic clusters, with *K* taking values of 1, 2, 3 and 4.

We investigated the ability of STRUCTURE to clarify introduction routes by focusing on *K* = 2 analyses. With *K* = 2, the two samples from an introduced population may or may not cluster together. With the scenario simulated here, in which the two invasive populations result from two independent introductions, the two samples of the introduced populations would not be expected to cluster together (Fig. 1). Indeed, the two independent drift pulses at work during these two introductions (i.e. the bottleneck events) should make the introduced populations more genetically different from each other than from the native population, from which they are separated by a single drift pulse. Consequently, STRUCTURE would yield a misleading pattern if the native population sample belonged to one cluster and the two invasive population samples both belonged to the other at *K* = 2. Indeed, this could be considered evidence for a lack of independence of the two populations, with one invasive population being the source of the other (Fig. 1; Fig. 2b). Such a clustering pattern, hereafter referred to as “misleading clustering”, would lead most STRUCTURE users to an incorrect interpretation, according to which a “successive introductions” scenario would be more likely than the “independent introductions” scenario. Note that STRUCTURE analyses carried out on three population samples with *K* = 3 are, theoretically, unsuitable for comparisons of independent and successive introduction scenarios, because each population sample would probably form its own cluster (Fig. 1).

For analysis of the 225,000 STRUCTURE runs with *K* = 2 and estimation of the frequency of misleading clusterings, the STRUCTURE output was characterized as follows. From the output file of each run, we extracted the proportion of membership *Qi_A_* and *Qi_B_* of population sample *i* for clusters *A* and *B*, respectively (with *Qi_B_* = 1 - *Qi_A_*). The *Qi_A_* and *Qi_B_* values were coded as *0*, *25*, *50*, *75* or *100* when belonging to the [0;0.2], ]0.2;0.4[, [0.4;0.6], ]0.6;0.8[ or [0.8;1] intervals, respectively. For each STRUCTURE run, we summarized the clustering pattern by a code *C*_1A_/*C*_2A_/*C*_3A_, where *C*_iA_ is the membership code of population sample *i* for cluster *A.* For example, the clustering code would be *0/0/100* for a STRUCTURE run output in which *Q1_A_* = 0.12, *Q2_A_* = 0.05 and *Q3_A_* = 0.96. Note that belonging to cluster *A* or *B* has no specific meaning, and the subscripts *A* and *B* can thus be permuted. For example, clustering codes *0/0/100* and *100/100/0* summarize the same pattern and are pooled together as *0/0/100.* Given the simulated scenario of independent introductions of the two invasive populations, *0/100/100* was the code considered to correspond to misleading clustering (Fig. 2b). All other clustering codes were considered non-misleading in the context of introduction routes inference. Focusing on the codes instead of the proportions of membership made it possible to pool together slightly different clustering patterns in the same category.

Given the stochastic processes involved in the MCMC analysis, the ten replicated STRUCTURE runs performed on a single dataset could conceivably generate different results, a phenomenon called genuine multimodality (Jakobsson and Rosenberg, 2007; Porras-Hurtado *et al.*, 2013). Clustering results for a given dataset were considered to be homogeneous if the same clustering code (as defined above) was obtained in all ten runs. They were otherwise considered to be heterogeneous. We evaluated the global occurrence of misleading clustering in the analyses of the simulated datasets, and focused on two critical categories of misleading clusterings (Fig. 2c):

i. “Misleading homogeneous clusterings”: for one dataset, all ten runs homogeneously provide the misleading clustering pattern *0/100/100.*
ii. “Misleading heterogeneous clusterings”: for one dataset, the ten runs are not homogeneous (i.e. genuine multimodality is observed) and the misleading clustering pattern *0/100/100* predominates.

For each dataset, we also inferred the best value of *K,* as follows: if the mean natural logarithm of the likelihood of the data *ln(P*(*X*|*K*)) with *K* in [1, 2, 3, 4] is maximal for *K* = 1, then the inferred number of clusters is 1; otherwise, we determined the best value of *K* (either *K* = 2 or *K* = 3) by the Δ*K* method (Evanno *et al.*, 2005).

### Effect of demographic parameter values on misleading clustering

For each number of simulated microsatellite loci (10, 20 or 100), the variables “proportion of analyses yielding misleading homogeneous clusterings” and “proportion of analyses yielding misleading heterogeneous clusterings” were analyzed independently with a generalized linear model, using a binomial probability distribution of the residual error and a logit link function. The following factors were included as fixed effects: effective population size *N* and bottleneck severity log_10_(*N*/*NF*). We used the Akaike information criterion (AIC) to select the best model from the various models of different complexity. Analyses were performed with R software V3.2.2 (R Development Core Team, 2015).

### Link between summary statistics of genetic diversity and STRUCTURE patterns

We summarized each simulated dataset, by using ARLSUMSTAT version 3.5 software (Excoffier and Lischer, 2010) to compute the mean number of alleles and the mean expected heterozygosity in each population sample, and the pairwise *F*_ST_ values between each pair of populations. We also used in-house PERL scripts to compute (i) the mean individual assignment likelihood (Rannala and Mountain, 1997) (*L*_*i*→j_) of each invading population (samples 2 and 3) to each possible source population (i.e. either the native population or the other invasive population), and (ii) the number of alleles shared by the invasive population samples.

For the comparison of datasets leading to “misleading homogeneous clusterings”, “misleading heterogeneous clusterings” and “non-misleading clusterings”, we specifically explored a few genetic diversity summary statistics: (i) expected heterozygosity of the native population sample, (ii) mean expected heterozygosity of both invasive population samples and (iii) the ratio of alleles shared by the two invasive population samples to the total number of alleles in the two samples. For each summary statistic and each number of loci, pairwise Mann-Whitney tests with Holmes correction for multiple comparisons were performed.

We also compared STRUCTURE results with those obtained by two other methods traditionally used to identify source populations: (i) the “*F*_ST_-based method” and the (ii) the “assignment likelihood-based method” (Genton *et al.*, 2005; Pascual *et al.*, 2007; Ciosi *et al.*, 2008; Tepolt *et al.*, 2009; Thibault *et al.*, 2009; Papura *et al.*, 2012; Mallez *et al.*, 2015; Dieni *et al.*, 2016). For an “independent introductions” scenario, we would expect the *F*_ST_ between the two invasive population samples to be larger than the *F*_ST_ values between the native population and each of the invasive population samples (i.e. *F*_ST_ 2-3 > *F*_ST_ 1-2 and *F*_ST_ 2-3 > *F*_ST_ 1-3). We would also expect both invasive population samples to be best assigned to the native population sample (i.e. *L*_2→1_ > *L*_2→3_ and *L*_3→1_ > *L*_3→2_).

For each dataset, a global exact test for population genotypic differentiation (Raymond & Rousset, 1995a) was carried out with GENEPOP software version 4.3 (Raymond & Rousset, 1995b). If a dataset displayed no population differentiation, we made the prudent and standard decision of not trying to infer any evolutionary relationship between the population samples. Consequently, such datasets were considered to generate non-misleading results for all methods.

## Results

### Effect of demographic parameter values on simulated datasets

The 500 simulated datasets for each parameter set are summarized with some common statistics in Table S1. Decreasing effective population sizes (*N*) generate lower intra-population and higher inter-population genetic diversities. Increasing bottleneck severity (log_10_(*N*/*NF*)) generates lower intra-population genetic diversities for both invasive samples, and overall higher inter-population genetic diversity. The main impact of a larger number of loci is a decrease in the variance of all summary statistics. Overall, the chosen parameter values (for *N* and *NF*) yield a large number of different combinations of genetic diversity for evaluation of the ability of STRUCTURE software to explore introduction routes in different situations.

### Overall STRUCTURE results

The best value of *K* inferred was most frequently three (Fig. 3 and Fig. S1). The proportion of datasets for which the best number of clusters was *K* = 3 increased strongly with increasing numbers of loci (41.9%, 50.6% and 74.9% for 10, 20 and 100 loci, respectively). More than 80% of the simulated datasets for which *K* = 3 was inferred by the Δ*K* method had heterogeneous clustering codes (i.e. genuine multimodality) at *K* = 2 (Fig. 3). By contrast, when the number of inferred clusters was one or two, multimodality at *K* = 2 was found in less than 10% of all datasets.

**Figure 3:**
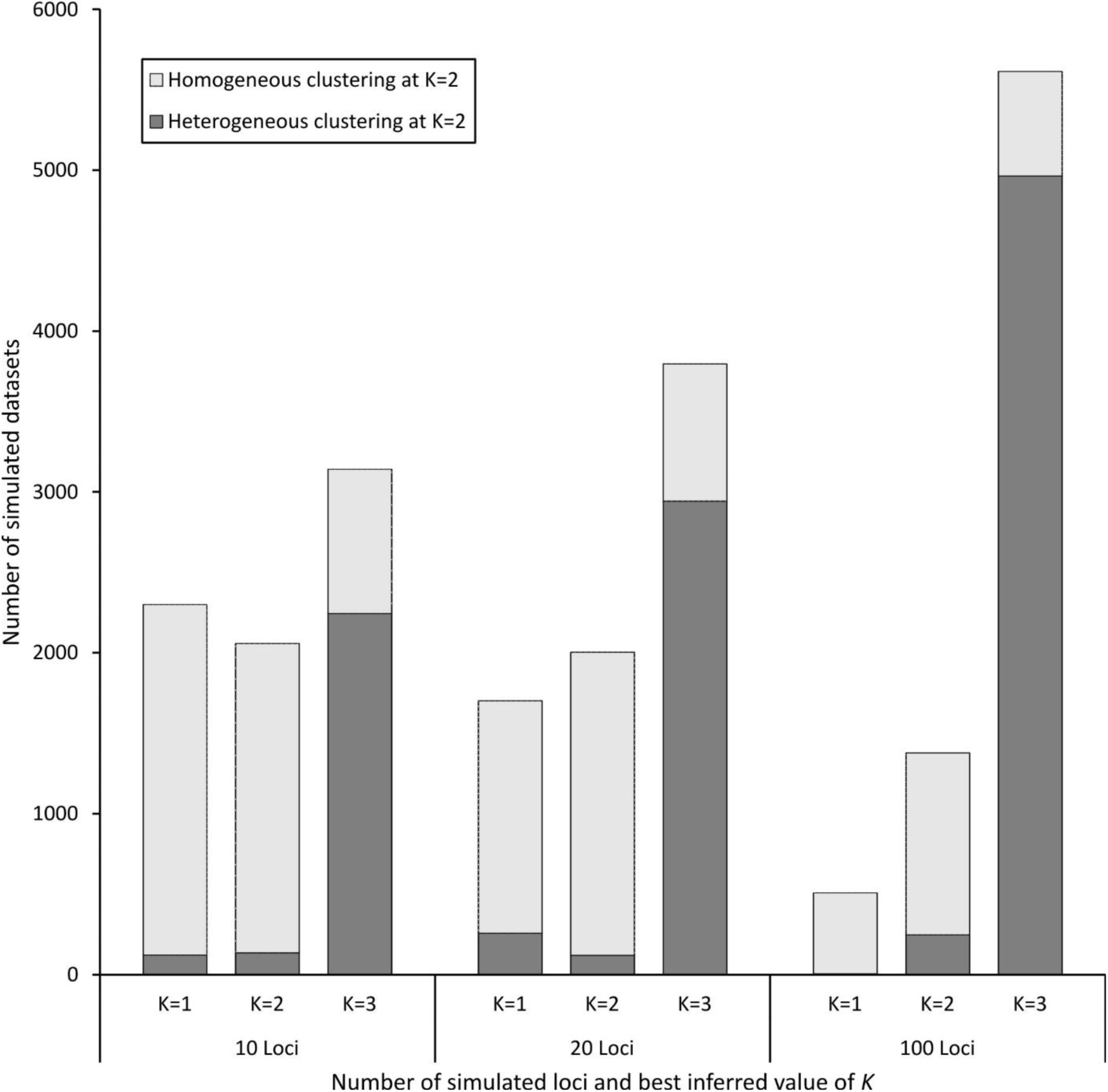
Distribution of the best number of clusters *K* inferred by Evanno ‘s method for each number of loci, and the proportion for which there was an absence (homogeneous clustering) or presence (heterogeneous clustering) of genuine multimodality in the ten STRUCTURE runs carried out at *K* = 2.

### Occurrence of misleading STRUCTURE patterns

Three categories of clustering codes at *K* = 2 accounted for more than 95% of all runs (see Table S2 for details): (i) clusterings in which all populations were fully admixed and undistinguishable with STRUCTURE (i.e. the *50/50/50* code), (ii) clusterings in which the two invasive samples belonged to different clusters (i.e. the *C*_1A_*/100/0* and *C*_1A_*/0/100* codes) and (iii) the misleading clusterings defined earlier (see Methods), in which the two invasive samples belonged to the same cluster, whereas the native sample belong to the other cluster (i.e. the *0/100/100* code, Fig. 2b).

Overall, the proportion of datasets with at least one misleading clustering pattern over the ten STRUCTURE runs (“misleading homogeneous clusterings”, “misleading heterogeneous clusterings” and non-misleading clusterings with at least one run yielding a misleading pattern) was 15.31%, 22.07% and 47.01% for 10, 20 and 100 simulated loci, respectively (Fig. 4a and Fig. S2), and very similar proportions were obtained with more (0.1 and 0.9) and less (0.3 and 0.7) stringent *Q_iA_* cutoff values (instead of 0.2 and 0.8 for *Q_iA_*) for the encoding of pattern results (Table S3).

**Figure 4:**
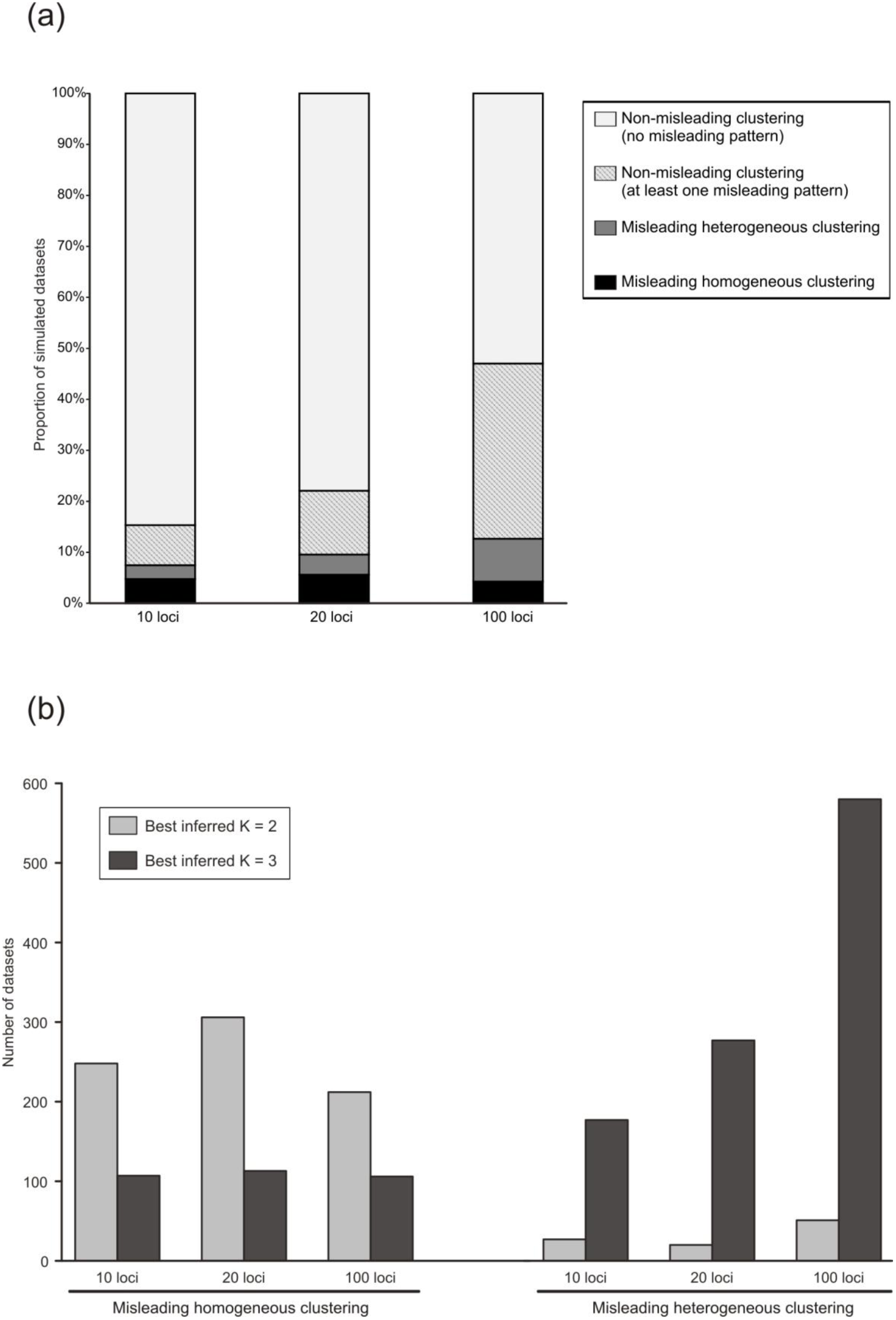
**(a)** Proportion of datasets with and without misleading patterns (Fig. 2b) for the ten STRUCTURE runs at *K = 2.* **(b)** Best inferred number of clusters K obtained by Evanno’s method for each number of loci within the datasets displaying misleading homogeneous (left) and misleading heterogeneous clusterings (right).

The frequency of “misleading homogeneous clusterings” was similar for different numbers of loci, and was rather low overall (between 4.24% and 5.59% of the datasets, Fig. 4a). “Misleading heterogeneous clusterings” were also infrequent, but their frequency increased with the number of loci: 2.71%, 3.96% and 8.41% for 10, 20 and 100 loci, respectively (Fig. 4a). Overall, 7.45%, 9.55% and 12.65% of datasets for 10, 20 and 100 loci, respectively, yielded misleading results. For some combinations of parameters, this proportion reached 36.8% of datasets (Fig. S2). *K* = 2 was most often (70%) inferred for datasets yielding “misleading homogeneous clusterings”, and *K* = 3 was most often (91%) inferred for datasets leading to “misleading heterogeneous clusterings” (Fig. 4b).

### Effect of demographic parameter values on STRUCTURE results

For the response variable “proportion of analyses yielding misleading homogeneous clusterings”, the best model according to the AIC always included the effective population size at equilibrium *N*, which was highly significant whatever the number of simulated loci (Table 1). Lower *N* values resulted in a higher proportion of misleading homogeneous clusterings (Fig. S3a). The best model also included bottleneck severity, log_10_(*N*/*NF*), and the interaction between the two main factors for 10 and 100 loci. Bottleneck severity was significant only for 10 loci, and had a positive effect: the stronger the bottleneck, the higher the proportion of misleading homogeneous clusterings. The interaction between the two factors was significant in both models (Table 1 and Fig. S3a).

**Table 1:** Results obtained with the best model selected from the various statistical models run for the response variables “proportion of analyses yielding homogeneous misleading clusterings” and “proportion of analyses yielding heterogeneous misleading clusterings”. Note: significant *P*-values, for a 5% threshold of significance, are shown in bold

| *Response variable* | *Number of loci* | *Factors of selected model* | *Estimate* | *Std error* | *z-value (df=7499)* | *P* |
| --- | --- | --- | --- | --- | --- | --- |
| Proportion of analyses yielding homogeneous misleading clusterings |  |  |  |  |  |  |
|  | 10 | *N* | -0.004 | 0.0005 | -7.931 | **<0.0001** |
| | | $\log_{10}(N/NF)$ | 0.399 | 0.084 | 4.746 | **<0.0001** |
| | | $N \times \log_{10}(N/NF)$ | 0.001 | 0.0002 | 5.185 | **<0.0001** |
|  | 20 | *N* | -0.005 | 0.0005 | -9.658 | **<0.0001** |
|  | 100 | *N* | -0.0003 | 0.00006 | -5.757 | **<0.0001** |
| | | $\log_{10}(N/NF)$ | -0.194 | 0.259 | -0.749 | 0.4540 |
| | | $N \times \log_{10}(N/NF)$ | -0.028 | 0.005 | -5.190 | **<0.0001** |
| Proportion of analyses yielding heterogeneous misleading clusterings |  |  |  |  |  |  |
|  | 10 | *N* | -0.0002 | 0.00007 | -3.496 | **0.0005** |
| | | $\log_{10}(N/NF)$ | 1.107 | 0.114 | 9.672 | **<0.0001** |
| | | $N \times \log_{10}(N/NF)$ | 0.00004 | 0.00002 | 1.827 | 0.0677 |
|  | 20 | *N* | -0.0004 | 0.00006 | -5.977 | **<0.0001** |
| | | $\log_{10}(N/NF)$ | 0.574 | 0.079 | 7.178 | **<0.0001** |
| | | $N \times \log_{10}(N/NF)$ | 0.00009 | 0.00002 | 4.854 | **<0.0001** |
|  | 100 | *N* | -0.0009 | 0.0001 | -9.666 | **<0.0001** |
| | | $\log_{10}(N/NF)$ | -0.801 | 0.068 | -11.715 | **<0.0001** |
| | | $N \times \log_{10}(N/NF)$ | 0.0003 | 0.00003 | 11.245 | **<0.0001** |

For the response variable “proportion of analyses yielding misleading heterogeneous clusterings”, the full model was selected for all numbers of simulated loci (Table 1). The effective population size at equilibrium *N* was significant in all cases, and had a negative effect (Fig. S3b). Bottleneck severity log_10_(*N*/*NF*) was also strongly significant for all numbers of loci, but its effect was positive for 10 and 20 loci and negative for 100 loci. The interaction between the two factors was significant for 20 and 100 loci, with a positive effect (Table 1).

### Links between summary statistics for genetic diversity and STRUCTURE patterns

The diversity of the native population, as assessed by its expected heterozygosity in the datasets with “misleading homogeneous clusterings”, was lower than that for “non-misleading” datasets, whatever the number of loci considered (Fig. 5). On the contrary, no clear trend could be observed for datasets with “misleading heterogeneous clusterings”. For these datasets, the mean expected heterozygosity was relatively high with 10 loci, intermediate with 20 loci and low with 100 loci, but, in each case, extreme low and high values were observed. The diversity of invasive populations, which was affected by both the diversity of the native population and bottleneck severity, was low for both kinds of misleading clusterings (Fig. 5). In comparisons with the “non-misleading” datasets, the proportion of alleles shared by the two invasive populations was higher for the datasets with “misleading homogeneous clusterings”, and lower for “misleading heterogeneous clusterings”, unless 100 loci were considered (Fig. 5).

**Figure 5:**
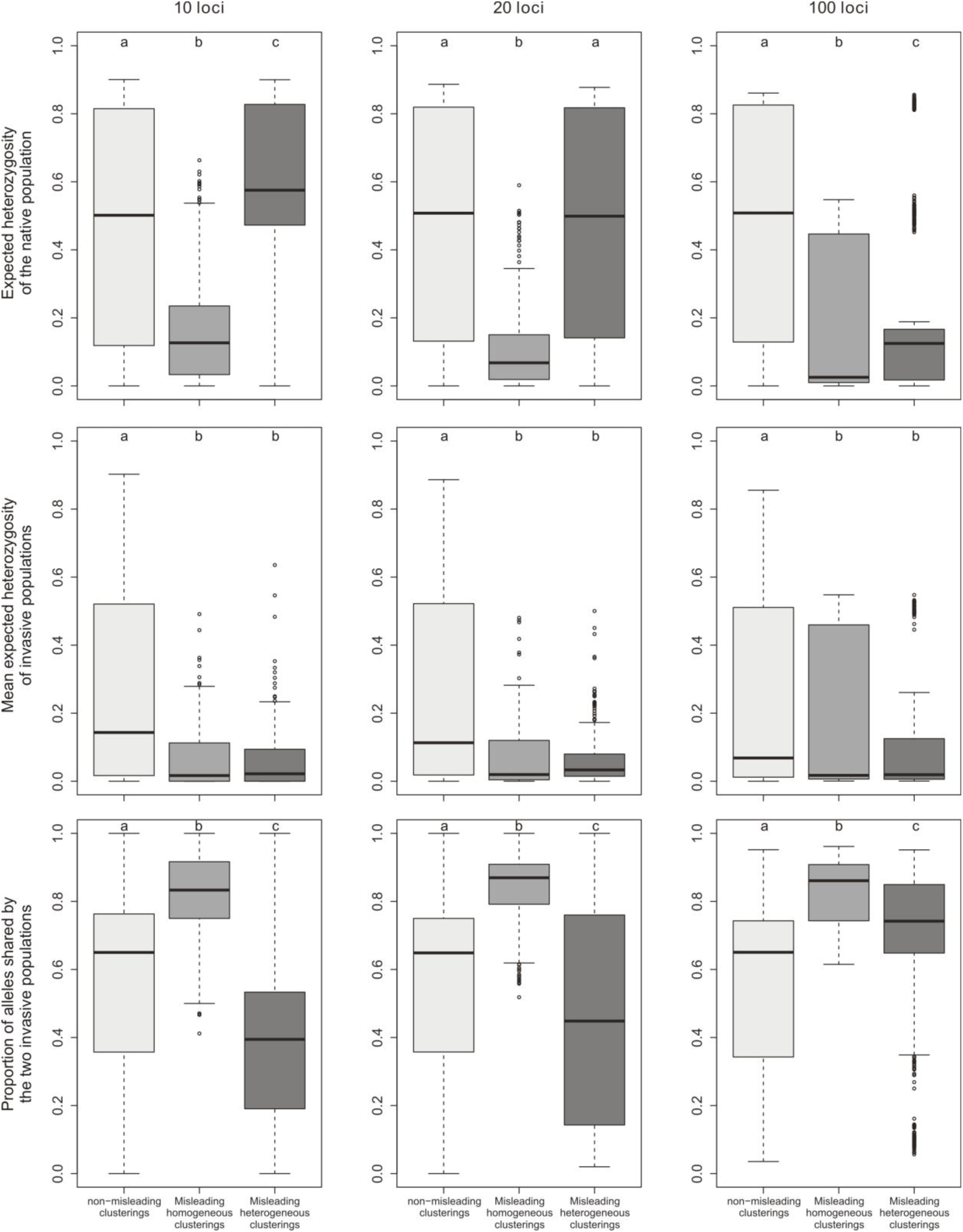
Tukey boxplots representing population genetics summary statistics for simulated datasets yielding non-misleading clusterings, misleading homogeneous clusterings or misleading heterogeneous clusterings. Within each frame, plots labeled with different letters are significantly different at the 5% level of significance (Mann-Whitney tests).

Outcomes for comparisons of STRUCTURE clusterings with results from *F*_ST_-based and assignment likelihood-based methods were very mixed, depending on the type of “misleading clusterings” considered. 86.5%, 93.6% and 99.1% of datasets with “misleading homogeneous clusterings” in STRUCTURE provided misleading results with at least one of the methods based on *F*_ST_ or assignment likelihood, when considering 10, 20 and 100 loci, respectively (Fig. S4). By contrast, datasets with “misleading heterogeneous clusterings” in STRUCTURE analysis were rarely (for 10 and 20 loci), or at least not as strongly (for 100 loci), associated with misleading results with the other methods: this was the case for 15.2%, 24.9% and 67.5% of these datasets for 10, 20 and 100 loci, respectively (Fig. S4). Note that, overall, STRUCTURE generates less misleading results than the other two methods.

## Discussion

We used simulated microsatellite datasets for a particular invasion scenario to determine whether the method implemented in the widely used STRUCTURE software (Pritchard *et al.*, 2000) could mislead users trying to infer introduction routes. We focused on a scenario with two independent introductions from a native population because this scenario can be rejected when obtaining some particular clustering results, which is not true for successive introductions scenarios when the chronology of introductions is not known. We found that, for a true scenario of two independent invasions from a single source, STRUCTURE runs could give misleading clustering patterns (i.e. the two invasive populations clustered together at *K* = 2). In about 10% of all simulated datasets, the results led to incorrect interpretation, with all (“homogeneous misleading clusterings”) or most (“heterogeneous misleading clusterings”) of the runs for a given dataset yielding the misleading pattern. Some combinations of demographic parameters resulted in higher frequencies of misleading results with STRUCTURE, and, contrary to expectations, increasing the number of loci also lead to an overall increase in the frequency of misleading results. Our results suggested that the two types of misleading clustering hazard, homogeneous and heterogeneous misleading clusterings, were very different. We suggest that (i) “homogeneous misleading clusterings” probably arise from a large probability of independently drawing the same alleles twice from a native population with low genetic diversity and that (ii) “heterogeneous misleading clusterings” probably randomly arise from convergence problems in STRUCTURE.

For “homogeneous misleading clustering”, the effective size of the native population has the strongest effect: the smaller this effective population size, the higher the risk of obtaining misleading clustering patterns over all STRUCTURE runs. Such “homogeneous misleading clustering” occurred principally when the two invasive populations shared a large proportion of alleles, and the *F*_ST_-based and likelihood assignment-based methods frequently yielded the same clustering pattern. Accordingly, the number of clusters inferred by the Evanno’s method was most frequently *K* = 2. Invasive populations encounter founder effects and genetic drift (Simberloff, 2009; Lawson Handley *et al.*, 2011), which are random processes. The probability of independently drawing the same alleles twice, with similar frequencies, from a given native population is usually low (when random processes are at work), but can actually be quite large when the diversity of the native population is itself low. This is particularly true in cases of low heterozygosity (Allendorf, 1986), in which one or a few alleles occur at high frequency at each locus.

The interpretation of “heterogeneous misleading clusterings” is less clear-cut, but several lines of evidence suggest the involvement of convergence issues in STRUCTURE runs. Indeed, “heterogeneous misleading clusterings” at *K* = 2 most often occurred when the best *K* value was undoubtedly 3, which corresponds to the true number of population. More generally, this category of misleading clusterings was associated with a better ability to differentiate the three populations. This may explain why the proportion of “heterogeneous misleading clusterings” was higher for a larger number of loci, for which more information is available to properly differentiate populations (Evanno *et al.*, 2005; Waples and Gaggiotti, 2006; Hubisz *et al.*, 2009). Besides, for 10 and 20 simulated loci, the proportion of “heterogeneous misleading clusterings” was positively related to bottleneck severity, which accentuates differences between populations. Overall, we suggest that “heterogeneous misleading clustering” probably results from a convergence problem in the MCMC procedure of STRUCTURE: when an inappropriate number of clusters is imposed — here *K* = 2 whereas the data are more consistent with *K* = 3 — multimodalities are often observed (Pritchard *et al.*, 2000; Jakobsson and Rosenberg, 2007), and sometimes, by chance, a large proportion of misleading clustering events occur in the various runs, resulting in “heterogeneous misleading clustering”.

### Conclusion and general recommendations

This study was based on a single simple invasion scenario with only three populations. More complex scenarios should be studied in the future, but this study constitutes a crucial first step, providing important information about the use of clustering methods in the context of biological invasions.

We found that STRUCTURE yielded misleading results, but at a low frequency. However, our results suggest that some situations should be analyzed with care. First, invasion biologists should be very cautious if the diversity of the native population is low: independent introductions from a single source population with low genetic diversity are likely to produce genetic signals similar to that expected for successive introductions. Such misleading results are difficult to spot, as they are consistent with the results of other methods, such as *F*_ST_- or assignment likelihood-based methods. In this context, quantitative methods, such as approximate Bayesian computation, may be very useful. Second, multimodal STRUCTURE results should be interpreted very cautiously, particularly if large numbers of loci are used. This is sobering news, because many published studies interpret STRUCTURE results at different *K* values, including those displaying genuine multimodality. This problem is not specific to the exploration of introduction routes and has much wider implications (Meirmans, 2015). Multimodality is often a sign of poor convergence of STRUCTURE runs, and is therefore likely to lead to results of limited biological meaning. In such situations, other methods (e.g. *F*_ST_-based, assignment likelihood-based) may make it possible to determine whether the STRUCTURE results are misleading or not. More generally, it is important to keep in mind that STRUCTURE results have to be interpreted cautiously (Pritchard *et al.*, 2010) and, in the context of invasion routes inferences, it should rather be used as a tool to clarify the scenery and decrease the number of genetic units from a large number of population samples to a few main clusters before quantitative analyses, such as approximate Bayesian computation, are performed (Lombaert *et al.*, 2014).

## Acknowledgments

We thank Margarita Correa, Arnaud Estoup, Thibaut Malausa and Ferran Palero for fruitful comments and discussions. We also thank Alexandre Dehne-Garcia for assistance with the computer cluster.

## Author contributions

E.L., T.G. and E.D. conceived and designed the study. E.D. wrote the scripts and ran the simulations and analyses. E.L. and E.D. analyzed the data. E.L., T.G. and E.D. wrote the paper.

